# Analysis of *Amblyomma americanum* microRNAs in response to *Ehrlichia chaff*eensis infection and their potential role in vectorial capacity

**DOI:** 10.1101/2024.05.03.592465

**Authors:** Deepak Kumar, Khemraj Budachetri, Yasuko Rikihisa, Shahid Karim

## Abstract

**Background:** MicroRNAs (miRNAs) represent a subset of small noncoding RNAs and carry tremendous potential for regulating gene expression at the post-transcriptional level. They play pivotal roles in distinct cellular mechanisms including inhibition of bacterial, parasitic, and viral infections via immune response pathways. Intriguingly, pathogens have developed strategies to manipulate the host’s miRNA profile, fostering environments conducive to successful infection. Therefore, changes in an arthropod host’s miRNA profile in response to pathogen invasion could be critical in understanding host-pathogen dynamics. Additionally, this area of study could provide insights into discovering new targets for disease control and prevention. The main objective of the present study is to investigate the functional role of differentially expressed miRNAs upon *Ehrlichia chaf*feensis, a tick-borne pathogen, infection in tick vector, *Amblyomma americanum*.

**Methods:** Small RNA libraries from uninfected and *E. chaffeensis*-infected *Am. americanum* midgut and salivary gland tissues were prepared using the Illumina Truseq kit. Small RNA sequencing data was analyzed using miRDeep2 and sRNAtoolbox to identify novel and known miRNAs. The differentially expressed miRNAs were validated using a quantitative PCR assay. Furthermore, a miRNA inhibitor approach was used to determine the functional role of selected miRNA candidates.

**Results:** The sequencing of small RNA libraries generated >147 million raw reads in all four libraries and identified a total of >250 miRNAs across the four libraries. We identified 23 and 14 differentially expressed miRNAs in salivary glands, and midgut tissues infected with *E. chaffeensis*, respectively. Three differentially expressed miRNAs (miR-87, miR-750, and miR-275) were further characterized to determine their roles in pathogen infection. Inhibition of target miRNAs significantly decreased the *E. chaffeensis* load in tick tissues, which warrants more in-depth mechanistic studies.

**Conclusions:** The current study identified known and novel miRNAs and suggests that interfering with these miRNAs may impact the vectorial capacity of ticks to harbor *Ehrlichia*. This study identified several new miRNAs for future analysis of their functions in tick biology and tick-pathogen interaction studies.

## Introduction

MicroRNAs (miRNAs) are non-coding RNAs with a size ranging from 18-25 nucleotides and play a significant role in post-transcriptional gene regulation (Bartel, 2009; Bartel and Chen, 2004). Latest studies have revealed the significance of miRNAs in arthropod immunity and host-pathogen interactions (Momen-Heravi and Bala, 2018; Miesen et al., 2016). In animals, miRNAs regulate post-transcriptional gene expression by binding to the 3’-untranslated region (3’-UTR), but there are also instances where the miRNA binds to the 5’-untranslated regions (5’-UTR), or coding regions. Perfect complementarity of 2-8 nucleotides at the 5’ end of the miRNA (seed region) is necessary for miRNA regulation, and the remaining sequence of miRNA might carry mismatches or bulges (Bartel, 2009; Rigoutsos, 2009; Schnall-Levin et al. 2010). The miRNA is transcribed as a primary miRNA transcript and processed by Drosha and Pasha into a pre-miRNA. The pre-miRNA is exported to the cytoplasm and processed by Dicer into a mature miRNA, which is then loaded into the microRNA-induced silencing complex (miRISC) and targets the complementary mRNA for degradation (Asgari, 2018, Flynt et al., 2010). Small non-coding RNAs (SncRNAs), including miRNAs, have shown tremendous potential in gene regulation at the post-transcriptional level in animals, plants, and arthropods, including ticks (Bartel, 2004; Carrington et al., 2003; Lai, 2015; Griffiths-Jones et al., 2008). Although more than 800 tick species are present worldwide, ticks are underrepresented in available miRNA resources. Databases such as miRbase have 49 *Ixodes scapularis* miRNAs and 24 *Rhipicephalus microplus* miRNAs, while MirGeneDB 2.1 contains 64 *Ixodes scapularis* miRNAs (Fromm et al., 2022).

The lone-star tick (*Amblyomma Americanum)* is an aggressive human-biting tick species, a known vector of numerous disease-causing agents, including *Ehrlichia chaffeensis*, *E. ewingii*, heartland virus, Bourbon virus, *Francisella tularensis*, *Borrelia lonestari* (Sanchez-Vicente and Tokarz, 2023). Lone star tick bites are also known to cause a food allergy, Alpha-Gal Syndrome (AGS), or red meat allergy (Commins et al., 2011; Crispell et al., 2019; Sharma and Karim 2022; Sharma et al., 2024). *Am. americanum* ticks are prevalent in the southern United States and have expanded their geographic range to the northeastern United States, and Canada (Stafford et al., 2018; Nelder et al., 2019). *E. chaffeensis*, a tick-borne Gram-negative obligatory intracellular bacterium, causes a severe flu-like febrile disease called human monocytic ehrlichiosis (HME), a prevalent life-threatening disease (Adams et al., 2017). Ehrlichiosis is an underreported tick-borne disease, and the pathogen infection of *E. chaffeensis* within the tick vector is a black box, and dynamics of vectorial capacity are largely unknown.

Given the contribution of miRNAs in numerous cellular processes, including development, immunity, and pathogen response in arthropods, the functional characterization of tick miRNAs in tick biology, and host-pathogen interactions remains to be investigated (Alvarez-Garcia et al., 2005; Saldana et al., 2017). Several omics studies have characterized the time-dependent, tissues-dependent blood-meal and pathogen-induced differential gene expression in variety of tick species (Karim et al., 2011; Karim and Ribeiro 2015; Guizzo et al., 2022; Adegoke et al., 2023; 2024; Anderson et al., 2008; Villar et al., 2015; Antunes et al., 2019; Popara et al., 2015; Bartel, 2004). However, studies addressing the role of pathogens in differentially modulating tick’s small RNAs are limited (Kumar et al., 2022; Artigas-Jerónimo et al., 2019; Hermance et al., 2019; Ramasamy et al., 2020). A handful of studies have investigated how miRNAs regulate the tick-pathogen interaction. These studies have shown that tick miRNAs promote the transmission of *Anaplasma phagocytophilum* and Powassan virus, thereby facilitating infection establishment in *Ixodes scapularis* (Artigas-Jerónimo et al., 2019; Hermance et al., 2019; Ramasamy et al., 2020). An elegant study demonstrated the tick miRNA-mediated regulation of vertebrate host gene expression at the tick-host interface (Hackenberg et al., 2017). However, regarding miRNA-mediated gene expression, the lone star tick (*Am. americanum*) is an underrepresented tick species, and there is an urgent need to investigate how *E. chaffeensis* utilizes tick miRNAs to promote its survival and persistence within the tick vector. This knowledge gap is of great concern, especially considering the increasing threat of the lone star tick to public health significance.

Understanding the molecular interactions between tick vectors and *E. chaffeensis*, and the characterization of differentially regulated tick miRNAs are needed to develop new approaches to combat tick-borne infections. Identification of new and novel miRNAs by using new sequencing platforms opened up a new avenue of research (Kumar et al., 2022; Artigas-Jerónimo et al., 2019; Luo et al., 2022). In this work, a small RNA sequencing approach was utilized to identify miRNAs induced by *E. chaffeensis* in *A. americanum* tissues, and differentially expressed miRNAs were functionally characterized using a miRNA inhibitor approach.

## Materials and Methods

### Ethics statement

All animal experiments were performed in strict accordance with the recommendations in the NIH Guide for the Care and Use of Laboratory Animals. The Institutional Animal Care and Use Committee of the University of Southern Mississippi approved the protocol for blood feeding of field-collected ticks (protocol # 15101501.3).

### Ticks and tissue dissections

Adult *Am. americanum* ticks, both infected and uninfected were prepared as described (Karim et al., 2012). Briefly, lab-grown *E. chaffeensis* (Arkansas strain) was microinjected in engorged nymphs and 1x DMEM media was injected as a control cohort for uninfected ticks (Karim et al., 2012; Budachetri et al., 2020:2022; Adegoke et al., 2024a; 2024). The injected engorged nymphs were kept in sterile vials with a piece of filter paper. The engorged nymphs molted into either unfed male or female adult ticks within 2 months. The ticks were maintained under standard conditions as outlined by Patrick and Hair (1975). A qPCR assay was used to determine the infection level of *E. chaffeensis* in freshly molted ticks (Dunphy et al., 2014). Uninfected and *E. chaffeensis* adult ticks were infested on a rabbit for blood-feeding (see the experimental design Figure 1). Partially blood-fed uninfected and pathogen-infected female ticks were removed from the rabbit, and tick tissues were dissected as described earlier (Karim et al., 2012), and dissected tissues were directly stored in Trizol (Life Technologies, Carlsbad, CA, USA). Samples were kept at −80°C until use.

**Figure 1.**
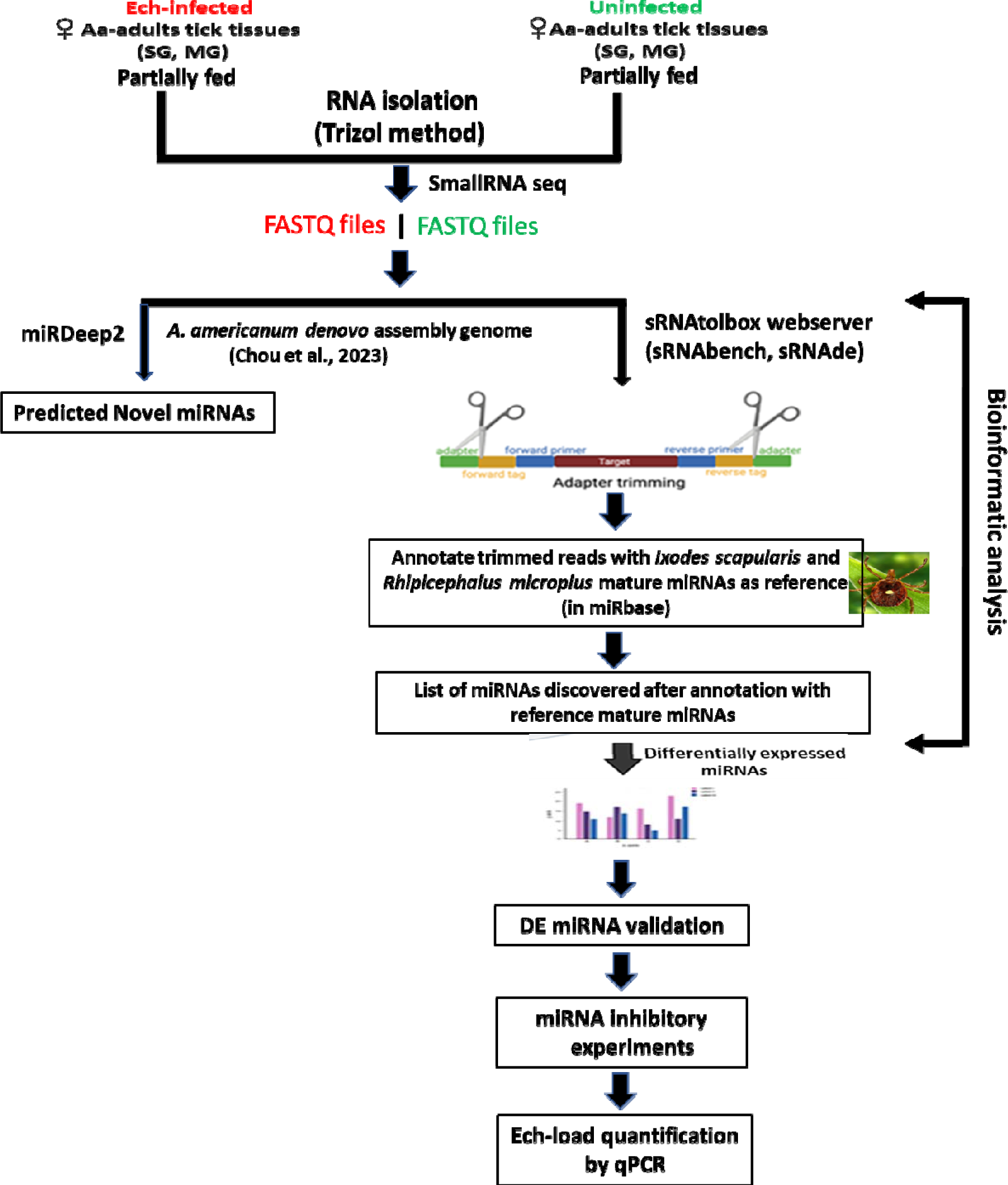
Schematic workflow of microRNA study in lone star tick tissues.

### RNA extraction and small RNA sequencing

RNA was extracted using Trizol RNA extraction methods from ten pooled tick tissue samples (uninfected, *E. chaffeensis*-infected midguts, and salivary glands). The integrity of the extracted RNA was determined using the standard spectrophotometric method as described earlier (Kumar et al., 2022). Small RNA library synthesis and small RNA sequencing were outsourced to the University of Mississippi Medical Center (UMMC) Molecular and Genomics Core laboratory. Briefly, four small RNA libraries (uninfected midguts, *E. chaffeensis* infected midguts, uninfected salivary glands, and *E. chaffeensis* infected salivary glands) were prepared using the Illumina Truseq Kit according to the manufacturer’s guidelines. Short adapter oligonucleotides were ligated to each end of the small RNAs. Following this, a cDNA copy was made with reverse transcriptase, and polymerase chain reaction (PCR) was utilized to incorporate sample-specific barcodes and Illumina sequencing adapters. The final concentration of all Next Generation Sequencing (NGS) libraries was determined with a Qubit fluorometric assay and a DNA 1000 high-sensitivity chip on an Agilent 2100 Bioanalyzer was used to assess the DNA fragment size for each library followed a purification step by polyacrylamide gel-electrophoresis. The sample libraries were pooled and sequenced on an Illumina Next Seq 500 (single end 36 bases) using TruSeq SBS kit v3 (Illumina) according to the manufacturer’s protocols.

### Data Analysis

There are currently two computational strategies available for miRNA profiling: 1) the genome-based strategy, which maps small RNA-seq reads to a reference genome and evaluates sequences that generate the characteristic hairpin structure of miRNA precursors (Bortolomeazzi et al., 2019), and 2) the machine-learning-based strategy, where biogenesis features of sequences are extracted based on the available miRNA sequences in microRNA databases such as miRBase (Kozomara et al., 2014) and from the analysis of miRNA duplex structures (Vitsios et al., 2017). During the analysis of this data, the *Amblyomma americanum* genome sequence was not available. Therefore, microRNA data was analyzed by the smallRNAtoolbox webserver (Aparicio-Puerta et al., 2022). Recently, Chou et al., (2023) sequenced the genome of the *Am. americanum* assembled the long-read sequenced genome and provided a file of a partially annotated genome. The miRDeep2 software package version 2.0.0.8 (Friedländer et al., 2012) was used to predict the novel and known miRNAs in all tick tissue samples using the partially annotated genome of *Am. americanum* (Supplementary Table S1-5). sRNAtoolbox algorithm has not been optimized for novel microRNA discovery or may have limitations in accurately identifying and characterizing novel microRNAs.Since the genome of the tick *Am. americanum* has not been fully annotated for noncoding RNAs such as rRNAs, tRNAs, snRNAs, snoRNAs, etc., the percentage abundance of these non-coding RNAs in our data has not been determined.

### Bioinformatics

As mentioned above, our small RNA data were analyzed by using a smallRNAtoolbox webserver (Aparicio-Puerta et al., 2022). Briefly, after an initial sequencing quality control step in FastQC (http://www.bioinformatics.babraham.ac.uk/projects/fastqc), preprocessing, mapping, and annotation were mainly conducted in sRNAbench (Aparicio-Puerta et al., 2022) with customized scripts as necessary. Simply, the obtained sequence reads were 36 nucleotides (nt) in length, but many small RNAs were between 27 and 33 nt. When forcing the detection of at least 10 nt of adapter as a typically used minimum length, only RNA molecules of up to 26 nt can be resolved (read length plus minimum adapter length). Therefore, to detect all small RNAs <36 nucleotides, we implemented iterative adapter detection and trimming. First, the adapter was detected in the whole read, and, if not found, it was then searched using iteratively shorter minimum adapter lengths at the 3′ end. After adapter trimming, the reads collapsed into unique reads followed by read count assignment, i.e., counting the number of times that each unique read was sequenced.

### Differential expression and normalization

Our experimental design resulted in several possible comparisons: (i) uninfected versus *E.chaffeensis* infected salivary glands (SGs), (ii) uninfected versus *E. chaffeensis* infected midgut (MGs), (iii) uninfected MGs versus uninfected SGs, and (iv) *E. chaffeensis*-infected MGs versus *E. chaffeensis*-infected SGs. An edgeR tool in the sRNAtool box was used to determine differential miRNAs expression between uninfected and *E. chaffeensis-*infected tick tissues (Robinson et al. 2010). Briefly, using the differential expression module of sRNAtoolbox (i.e. sRNAde), we generated an expression matrix with the raw read counts for input into edgeR to obtain differential miRNA expression. The edgeR normalizes the data using the trimmed mean of M-values (TMM) method. We also generated an expression matrix with reads per million (RPM)-normalized expression values using the “single assignment” procedure in sRNAbench. As a result, each read mapping multiple times was only assigned once to the miRNA with the highest expression and only affected reads mapping to several different reference sequences, i.e., normally miRNA sequences from the same family. The RPM values were obtained by dividing the read count of a given miRNA by the total number of reads mapped to the miRNA library.

### Validation of differentially expressed miRNAs

All miRNAs that were differentially expressed in small RNA sequencing data were validated by qRT-PCR in *Am. americanum* tick tissues. Mir-X miRNA qRT-PCR TB Green kit (Takara BIO, San Jose, CA, USA) was used for cDNA synthesis and miRNA expression analysis. This kit includes the Mir-X miRNA First-Strand Synthesis Kit, which transforms RNA into complementary DNA (cDNA) and enables the quantification of particular miRNA sequences through real-time PCR. Briefly, utilizing a single-tube method, RNA molecules undergo polyadenylation and reverse transcription by the action of poly(A) polymerase and SMART® MMLV Reverse Transcriptase, both components of the mRQ Enzyme Mix are supplied with the kit. Subsequently, real-time qPCR is conducted using the TB Green Advantage® qPCR Premix and mRQ 3’ Primer, in combination with miRNA-specific 5’ primers, to quantify specific miRNA expression. Primers used are listed in supplementary Table S6. Conditions used for qRT-PCR were initial denaturation of 95° C for 10 mins, then 40 cycles of 95° C for 5 secs, and 60° C for 20 secs.

### MicroRNA inhibition assay

We selected three microRNAs, including Aam-miR-87, Aam-miR-750, and Aam-miR-275 for further characterization. A miRNA inhibitor approach works by sterically blocking specific miRNA functions using an oligonucleotide that complements the mature miRNA target (Lennox et al., 2013). These inhibitors were designed and synthesized by Integrated DNA Technologies (IDT, Coralville, IA), which also synthesized non-target negative controls alongside the specific inhibitors. Three groups of *E. chaffeensis*-infected ticks were injected with miRNA inhibitors for each selected miRNA (Aam-miR-87, Aam-miR-750, and Aam-miR-275), while a fourth group was injected with non-target negative controls. Each group contained 25 female ticks. Using a published study (Ramasamy et al. 2020), we selected a 1.05 nanomoles dose for this assay. Briefly, 1.05 nanomoles of each microRNA inhibitor and negative control were injected into each female tick within their respective groups. The injected ticks were allowed a 48-hour recovery period along with non-treated males (25 females/15 males) in a laboratory incubator, maintained at a temperature range of 23 ± 2°C with a humidity level of 95%, and under a light cycle of 14 hours of light and 10 hours of darkness. After the recovery period of 48 hrs, all groups of ticks were infested on a sheep in separate stockinet cells to blood feed. The partially blood-fed ticks were removed from the sheep, and tick tissues (MGs, and SGs) were dissected within 4 hrs for downstream analysis. RNA was extracted from each tissue using the Trizol method as described previously (Kumar et al., 2022). In a qPCR assay, the *E. chaffeensis* infection in individual tick tissues was assessed using disulfide bond formation (dsb) gene primers (Dunphy et al., 2014). The level of miRNA inhibition was determined by using QRT-PCR primers designed by the miRprimer2 algorithm (Busk, 2014), in conjunction with the Mir-X™ miRNA qRT-PCR TB Green® Kit from TAKARA (San Jose, CA, USA). After confirming miRNA inhibition in tick tissues, the relative *E. chaffeensis* load compared to the non-target negative control was quantified in the respective tick tissues.

## Results and Discussion

### Profile characteristics of small RNA libraries

The small RNA sequencing yielded a total of >147 million raw reads, comprised of >32 million from uninfected midgut, >32million from the *E. chaffeensis-*infected midgut, >40million from uninfected salivary glands, and >41 million from the *E. chaffeensis-* infected salivary glands. Following the adapter trimming process and subsequent removal of short reads (those ≤20 nucleotides (nt) in length), the small RNA reads left for downstream analysis were >41 million from *E. chaffeensis-*infected samples and >36 million from uninfected samples. The distribution of read length presents an indication of the types of small RNAs present in both *E. chaffeensis-*infected and uninfected tick tissues (midgut and salivary gland). Sequencing stats including raw reads, adapter cleaned reads, reads in analysis, quality filter reads have been provided in supplementary Table S7, while processing stats of reads (in percentage) and raw miRNA summary (number of detected miRNAs and its precursors) have been provided correspondingly as supplementary Table S8 and S9. Both types of samples, *E. chaffeensis-*infected and uninfected, exhibited two main peaks at 22 nt (representing miRNAs/siRNAs) and 29 nt (representing piRNAs) (Figure 2). Notably, PIWI-interacting RNAs (piRNAs) constitute a class of small RNAs, which usually vary in size from 26 to 31 nucleotides (Iwasaki et al., 2015; Santos et al., 2023). The piRNAs associate with PIWI proteins, which belong to the Argonaute family of proteins and are active in the testes of mammals. These RNAs play a crucial role in germ cell and stem cell development in invertebrates. One of their primary functions is the silencing of transposable elements (TEs) to protect genomic integrity (Aravin et al., 2008; Brennecke et al., 2007; Brennecke et al., 2008).piRNAs are passed down from the female germline to progeny, ensuring the stability of the genome across generations. When males harboring a specific family of transposon elements (TEs) mate with females devoid of these elements, the females lack the necessary complementary piRNAs to defend their genome resulting in an overabundance of transposons. Such proliferation could potentially lead to sterility, a situation termed hybrid dysgenesis (Erwin et al., 2015). This concept of hybrid dysgenesis holds potential for tick control.

**Figure 2.**
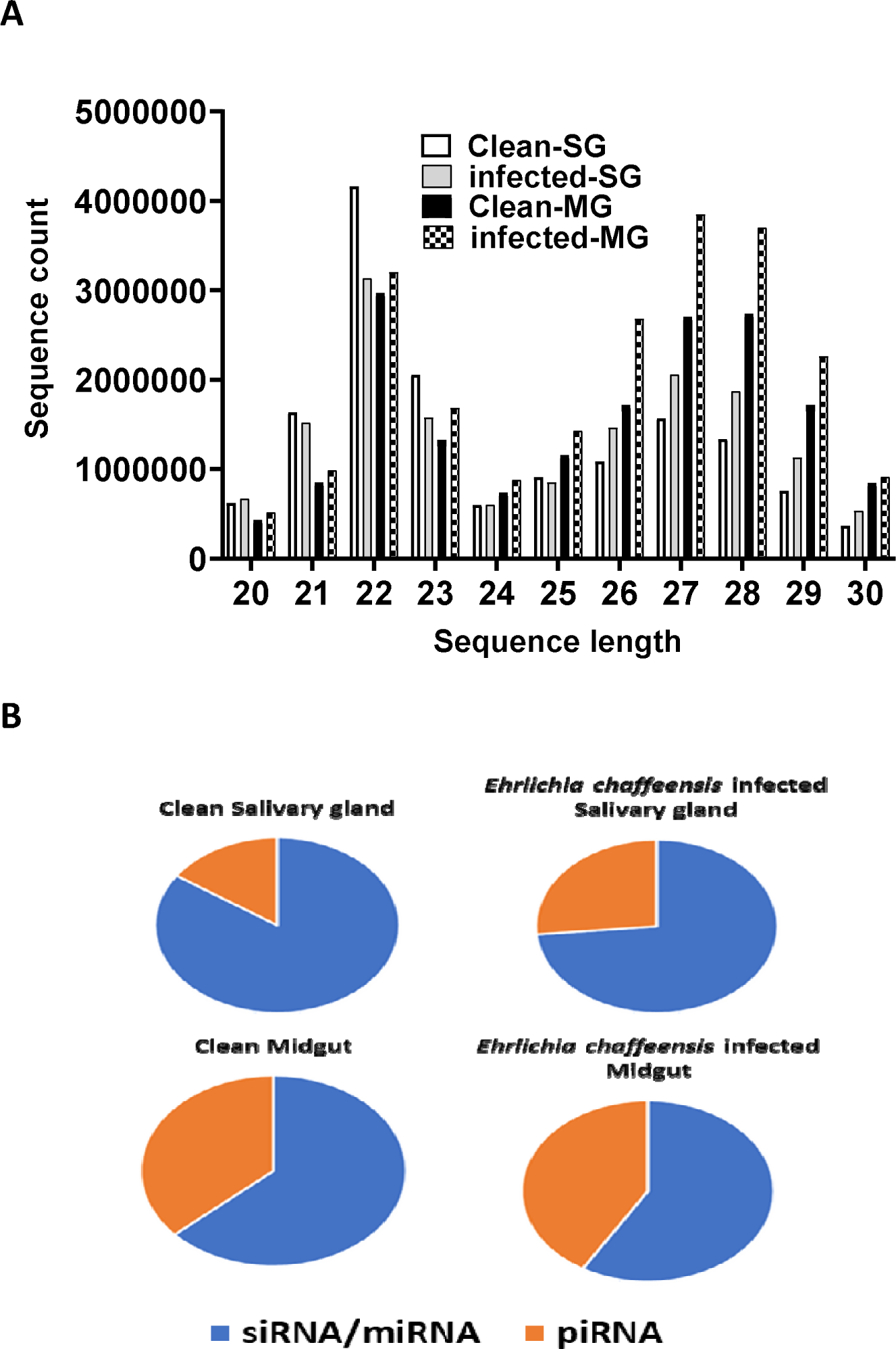
A. Small RNA sequence length distribution in uninfected (clean) and *Ehrlichia chaffeensis-*infected tick tissues (SG, MG). MicroRNAs are twenty-two (22) nucleotides in length. B. Pie-chart distribution of small RNA reads. There are mainly two characteristic peaks of tick small RNAs (miRNAs/siRNAs-22 nt, piRNAs-29 nt). Aa-*Amblyomma americanum*, PF-Partially fed (2 days), SG-Salivary glands, MG-Midgut, nt-nucleotide. Ech – *E. chaffeensis* infected.

### *In silico* mapping of *E. chaffeensis* infected small RNA sequences to *E. chaffeensis*Arkansas strain genome

Upon performing *in silico* mapping of small RNA reads infected with *E. chaffeensis* to the genome of the *E. chaffeensis* Arkansas strain (GCF_000013145.1_ASM1314v1_genomic.fna), we detected 1,413 *E. chaffeensis* sequences in the midgut out of a total of 2,492,851 reads. In addition, we found 3,185 *E. chaffeensis* sequences in the salivary glands from a total of 19,103,069 reads. From these results, it appears that our samples were infected with *E. chaffeensis*, and the infection level was low, a hallmark of Ehrlichia infection within the tick vector (Kennedy and Marshall, 2021).

### Differentially expressed microRNAs in *E. chaffeensis* infected tick salivary glands

Our small RNA sequencing analysis identified 360 microRNAs including known and predicted ones in uninfected and *E. chaffeensis-*infected tick tissues (Supplementary data Table S1). The list of differentially expressed miRNAs in *E. chaffeensis-*infected salivary glands (SGs) includes 18 upregulated miRNAs, Aam-miR-5322, Aam-miR-1, Aam-miR-750, Aam-miR-993, Aam-miR-5307, Aam-miR-87, Aam-miR-5315, Aam-miR-133, Aam-let-7, Aam-miR-3931, Aam-miR-263a, Aam-miR-8, Aam-miR-5305, Aam-bantam, Aam-miR-279, Aam-miR-5306, Aam-miR-276, Aam-miR-315) and 5 downregulated miRNAs Aam-miR-12, Aam-miR-2a, Aam-miR-10, Aam-miR-7, Aam-miR-285 (Figure 3).

**Figure 3.**
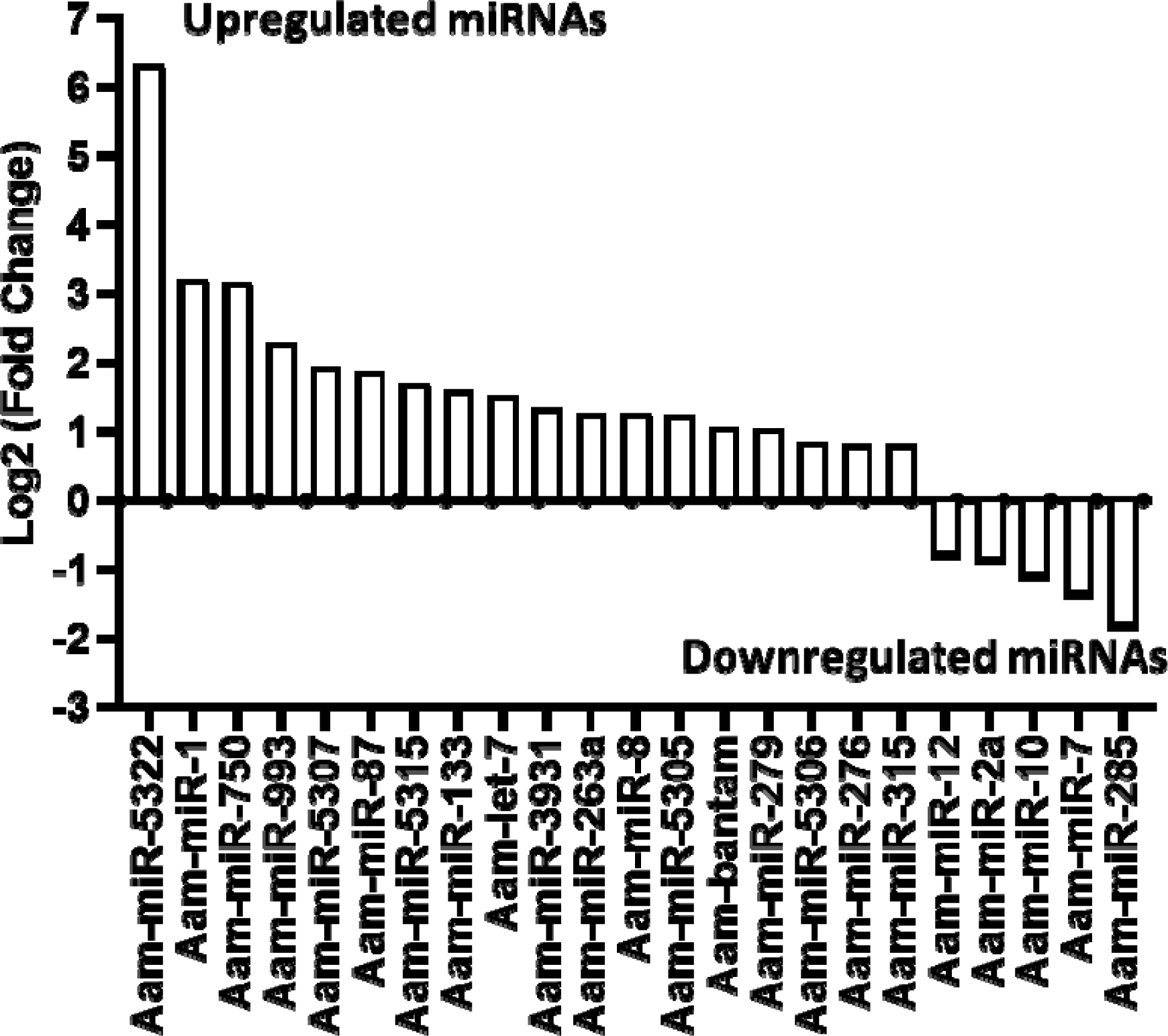
In silico differential expression of predicted microRNAs in *Ehrlichiachaffeensis*infected partially fed salivary gland relative to partially fed clean salivary glands. EdgeR was used for differential expression analysis. miRNAs with a Log2 fold-change expression > |0.8|, p-value < 0.05 were considered significantly differentially expressed.

### Differentially expressed microRNAs in *E. chaffeensis* infected tick midgut

Likewise, in the *E. chaffeensis* infected tick midgut (MG), seven miRNAs were upregulated in *Ehrlichia* infection, Aam-miR-5309, Aam-miR-79, Aam-miR-1, Aam-miR-275, Aam-miR-315, Aam-miR-278, Aam-miR-263a and 7 downregulated miRNAs, Aam-miR-153, Aam-miR-12, Aam-miR-7, Aam-miR-124, Aam-miR-5314, Aam-miR-5310, Aam-miR-285 (Figure 4). Heat map representation of DE miRNAs (Supplementary Figure S1-S2) has also been provided as supplementary information). All these miRNAs are listed in miRbase (22.1) and have also been identified in *Ixodes scapularis* or *Rhipicephalus microplus* tick species. It is necessary to investigate the roles of these differentially expressed miRNAs in *E. chaffeensis-*infected tick tissues. It is crucial to gain a deeper understanding through these differentially expressed miRNAs of how *E. chaffeensis* manipulates the expression of tick microRNAs to ensure its survival, persistence, and transmission. This knowledge holds immense potential in the development of innovative tools to effectively block the transmission of tick-borne pathogens.

**Figure 4.**
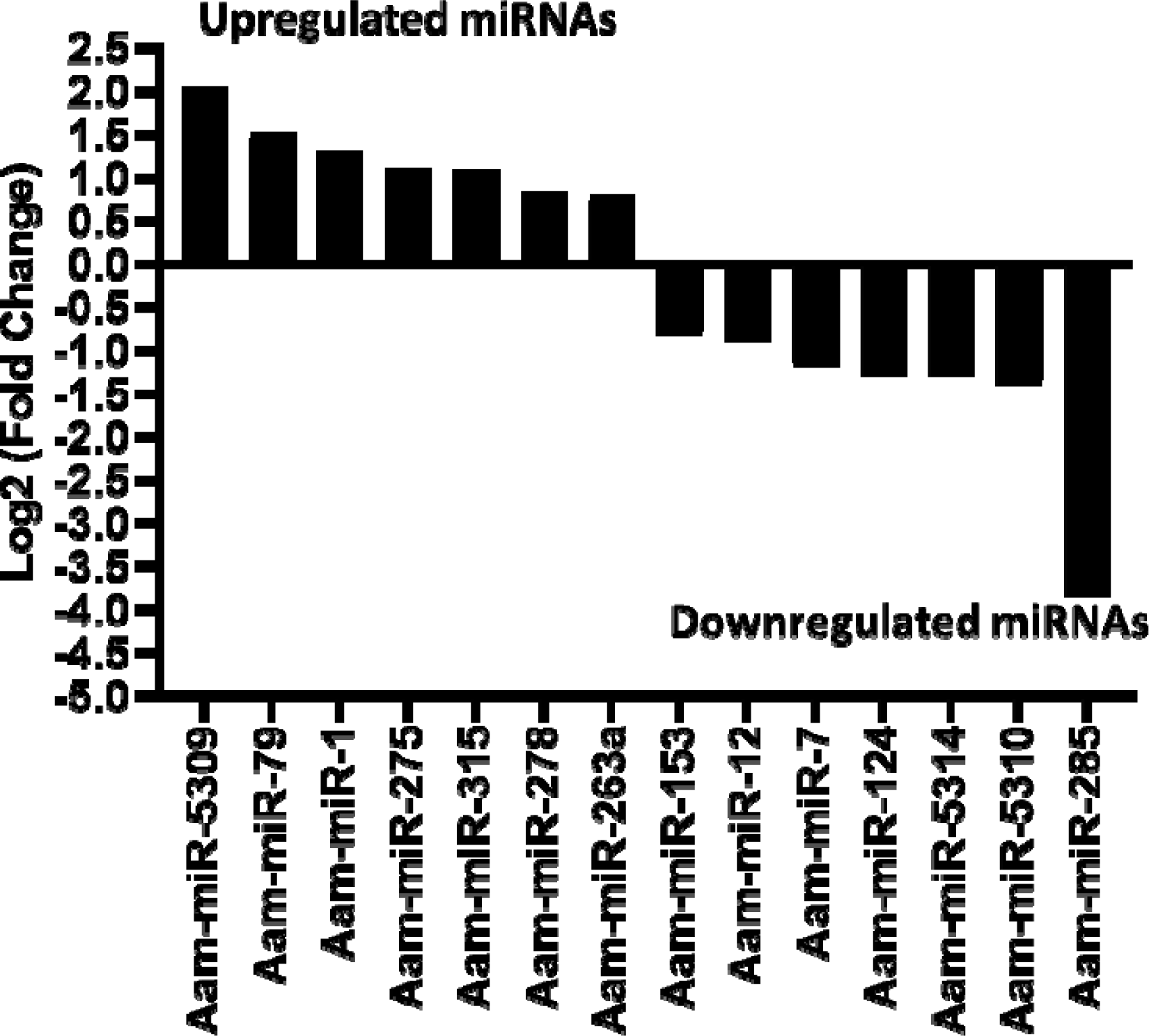
In silico differential expression of predicted microRNAs in *Ehrlichiachaffeensis* infected partially fed midgut relative to the partially fed clean midgut. EdgeR was used for differential expression analysis. miRNAs with a Log2 fold-change expression > |0.8|, p-value < 0.05 were considered significantly differentially expressed.

In this study, we found miR-1 upregulated in *E. chaffeensis* infected tick tissues, i.e. SG, and MG in comparison to uninfected ones. It is noteworthy that miR-1 often exhibits increased expression during pathogen infections. miR-1 belongs to a conserved family, which includes miR-7 and miR-34, and is conserved across various organisms such as fruit flies, shrimps, and humans (Takane et al. 2010). It participates in analogous pathways in these organisms, including development and apoptosis, and its upregulation has also been identified during stressful conditions (Huang et al., 2012). In mosquitoes, miR-1 is similarly upregulated during Plasmodium infection (Huang and Zhang, 2012), and facilitates West Nile Virus infection (Hussain et al., 2012). For *Bombyx mori*, the Nucleo-polyhedrosis virus (NPV) releases miR-1 inside the host to regulate its target RAN (exportin 5, co-factor), a player in miRNA biogenesis responsible for transporting pre-miRNA from the nucleus to the cytoplasm (Singh et al., 2012). Moreover, during *Listeria* infection in macrophages, miR-1 stimulates IFN-γ-dependent activation of the innate immune response (Xu et al., 2019). Based on these findings, we speculate that miR-1 could be upregulated in *E. chaffeensis-*infected tick tissues to trigger an immune response against *E. chaffeensis*.

The upregulation of miR-87 in *Ehrlichia*-infected SGs points out its putative role in ticks’ innate immune response. Earlier work on other arthropods, including *Manduca sexta* and *Aedes albopictus* hint at its potential role in interfering with innate immunity, particularly IMD and toll receptor signaling pathways (Zhang et al., 2014; Liu et al., 2015; Avila-Bonilla et al., 2017). Its *in silico* predicted targets in *Aedes albopictus* include TOLL pathway signaling Ser/Thr. Kinase, Toll-like receptor TOLL 1A, Class A Scavenger receptor with Se-Protease domain, and Galectin (Liu et al., 2015), as well as putative TLR 5b (Avila-Bonilla et al., 2017). In *Manduca sexta*, its predicted target is FADD, an adaptor protein involved in DISC formation (Zhang et al., 2014). Based on the published studies, we hypothesize that *Ehrlichia* infection differentially regulates the miR-87 to inhibit the Toll pathway for its survival in the tick vector.

The upregulation of miR-bantam in *Ehrlichia*-infected SGs also suggests its putative role in the pathogen infection of *E. chaffeensis*. Interestingly, miR-bantam, a conserved microRNA, exhibits high expression levels in insects. For example, it is highly expressed in *Drosophila* and participates in several key cellular processes such as cell proliferation, apoptosis, development, and the circadian rhythm. In *Drosophila*, miR-bantam serves two major roles: inhibiting apoptosis by down-regulating the apoptotic gene hid (Brennecke et al., 2003) and promoting cell proliferation by targeting genes like mad (Robins et al., 2005). The inhibition of apoptosis might act as a survival mechanism for *E.chaffeensis* within tick salivary gland cells. Our hypothesis warrants in-depth future studies to determine the role of miR-bantam in the vectorial capacity of the tick vector.

Another miRNA candidate, mir-79 was also significantly upregulated *Ehrlichia* infected MG. Earlier studies have described its role in various pathways, including immunity, cell differentiation, neurogenesis, and apoptosis. It has also been implicated in cancer and disease caused by viral infection (Artigas-Jerónimo et al., 2019; Yuva-Aydemir et al., 2011; Seddiki et al., 2013; Pedersen et al., 2013; Ouyang et al., 2015; Dong et al., 2017). It is known that mir-79 disrupts the JNK pathway by targeting its component genes *pvr* (CG8222) and *puc* (CG7850) (Fullaondo and Lee., 2012). The JNK pathway is an immune response pathway against Gram-negative bacterial pathogens (Bond and Foley, 2009). In the current study, mir-79 is upregulated in the *E. chaffeensis-infected* midgut. Similarly, in ticks infected with *Anaplasma phagocytophilum* (a Gram-negative bacterial pathogen), mir-79 was found upregulated, thereby facilitating infection by targeting the Roundabout protein 2 pathway (Robo2) (Artigas-Jerónimo et al., 2019). The upregulation of mir-79 could potentially be a mechanism used by *E. chaffeensis* to evade the tick’s immune system.

Our in-silico data reveals the downregulation of miR-5310 in the midgut of *E. chaffeensis-*infected ticks. MiR-5310, a tick-specific miRNA (Barrero et al., 2011), may play a role in tick feeding, as it was shown to be downregulated in Rhipicephalus microplus tick larvae exposed to host odor without being allowed to feed (Barrero et al., 2011). Another study showed its downregulation in *Anaplasma phagocytophilum*-infected nymphs compared to unfed, uninfected nymphs (Ramaswamy et al., 2020). Previous research has also shown the modulation of signaling events upon A. phagocytophilum infection of ticks (Khanal et al., 2018; Neelakanta et al., 2010; Sultana et al., 2010; Taank et al., 2017; Turck et al., 2019; Ramaswamy et al., 2020). Thus, in the case of the *E. chaffeensis-infected* tick midgut in the present study, miR-5310’s speculated role might be to modulate signaling events to protect *E. chaffeensis*. Its potential role could also be in tick feeding, as shown by its downregulation in *Rhipicephalus microplus* tick larvae exposed to host odor without being allowed to feed. In our data, miR-133 is upregulated in the salivary glands of *E. chaffeensis-infected* ticks. According to a recent study, the infection of ticks with the pathogen *Anaplasmaphagocytophilum* results in the downregulation of tick microRNA-133 (miR-133), leading to the induction of the Ixodes scapularis organic anion transporting polypeptide (isoatp4056) gene expression, which is critical for the pathogen’s survival within the vector and its transmission to the vertebrate host (Ramaswamy et al., 2020). Therefore, the upregulation of miR-133 in *E. chaffeensis-*infected tick salivary glands in our study might suggest the downregulation of organic anion transporting polypeptide (isoatp4056) gene expression, potentially inhibiting *E. chaffeensis* survival and transmission. This is a hypothetical explanation for the observed miR-133 upregulation, and further investigation is necessary for confirmation.

Let-7, an evolutionarily conserved microRNA in bilateral animals, plays a role in developmental regulation, such as molting and metamorphosis in arthropods, and can disrupt innate immunity by targeting the antimicrobial peptide diptericin (Carrington and Ambros, 2003; Hertel et al., 2012; Pasquinelli et al., 2000; Ling et al., 2014; Garbuzov and Tatar, 2010). Recent studies have also suggested its role in the molting of *Hyalomma asiaticum* ticks by targeting the ecdysteroid receptor (ECR), a part of the 20E signaling pathway (Sempere et al., 2003; Wu et al., 2019). In this study, let-7’s upregulation in tick salivary glands implies a possible role in targeting the antimicrobial peptide diptericin, potentially allowing *E. chaffeensis* to evade innate immunity. This may represent a mechanism for *E. chaffeensis*’s survival and successful transmission in tick salivary glands, but further investigation is required. It should be noted that diptericin inhibits Gram-negative bacteria by disrupting membrane integrity.

In this study, miR-275 is upregulated in the midgut of *E. chaffeensis-*infected ticks. MiR-275 directly targets and positively regulates the sarco/endoplasmic reticulum Ca2+ adenosine triphosphatase (SERCA), an active player in transporting Ca2+ from the cytosol to the sarco/endoplasmic reticulum (ER) in mosquito guts (Zhao et al., 2017). The transportation of Ca++ from the cytoplasm to the ER is required for the spreading process of *Ehrlichia canis* (Alves et al., 2014), suggesting that *E. chaffeensis* may modulate tick machinery via upregulation of miR-275. However, a follow-up study is necessary for confirmation. It’s also worth noting that miR-275 was found to be crucial for blood digestion and egg development in the mosquito *Aedes aegypti* (Bryant et al. 2010).

In our data set, miR-750 is upregulated in *E. chaffeensis*-infected tick salivary glands. Past studies have suggested its role in innate immunity, hormone signaling, and stress response (Rebijith et al., 2016; Nunes et al., 2013; Kanoksinwuttipong et al., 2022; Queiroz et al., 2020). A recent study indicated that upregulated miR-750 suppresses its target, the sarcoplasmic calcium-binding protein (Scp), and inhibits apoptosis, thus contributing to pathogen propagation (Kanoksinwuttipong et al., 2022). Given these previous studies, the possible role of miR-750 in inhibiting apoptosis and promoting *E. chaffeensis* propagation in tick salivary glands can be speculated. This could represent a mechanism by which *E. chaffeensis* avoids cellular apoptosis and propagates for effective transmission in the tick salivary glands. The roles of all the differentially expressed miRNAs mentioned above are listed in Table 1, along with other necessary details.

**Table 1.** List of differentially expressed microRNAs detected in *Ehrlichia chaffeensis* infected tick tissues (salivary gland, midgut) and their putative roles.

| microRNA | Role of ortholog microRNA | Target genes | Reference |
| --- | --- | --- | --- |
| miR-1 | Stress response, immunity, development, facilitating infection | HSP60, HSP70, GATA4, RAN (exportin) | Huang et al., 2012; Huang and Zhang, 2012, Hussain et al., 2012; Xu et al., 2019, Singh et al., 2012; Liu et al., 2017] |
| miR-79 | Innate immunity, differentiation, apoptosis | Roundabout protein 2 pathway (robo2), DRAPER, HP2 (Hemolymph protease), P38, *pvr*, *puc* | Fullaondo and Lee., 2012; Bond and Foley, 2009; Artigas-Jerónimo et al., 2019; Yuva-Aydemir et al., 2011; Seddiki et al., 2013; Pedersen et al., 2013; Ouyang et al., 2015; Dong et al., 2017 |
| let-7 | Developmental regulation such as molting and metamorphosis in arthropods, disrupts innate immunity | Antimicrobial peptide dipteracin | Pasquinelli et al., 2000; Ling et al., 2014; Garbuzov and Tatar, 2010; Sempere et al., 2003;Wu et al., 2019 |
| miR-133 | *Anaplasma phagocytophilum* survival inside ticks, transmission | Organic anion transporting polypeptide (isoatp4056) gene | Ramaswamy et al., 2020 |
| miR-bantam | Proliferation, apoptosis inhibition, development, circadian rhythm | *hid*, *mad* | Brennecke et al., 2003; Robins et al., 2005 |
| miR-87 | Pathogen survival (disruption of Toll and IMD pathway) | Serine/Threonine Kinase, Toll 1A, Putative TLR 5b, FADD | Zhang et al., 2014; Liu et al., 2015; Avila-Bonilla et al., 2017 |
| miR-5310 | Pathogen survival by modulating signaling pathways, Feeding behavior |  | Barrero et al., 2011; Khanal et al., 2018; Neelakanta et al., 2010; Sultana et al., 2010; Taank et al., 2017; Turck et al., 2019; Ramaswamy et al., 2020 |
| miR-275 | Active transport of Ca<sup>2+</sup> from cytosol to the endoplasmic reticulum, blood digestion, egg development |  | Zhao et al., 2017; Alves et al., 2014; Bryant et al., 2010 |
| miR-750 | innate immunity, hormone signaling and stress response, apoptosis inhibition | Sarcoplasmic calcium binding protein (Scp) | Rebijith et al., 2016; Nunes et al., 2013; Kanoksinwuttipong et al., 2022; Queiroz et al., 2020 |
| miR-153 | Development, immune response |  | Wu et al., 2013 |
| miR-5307 | Propagation of Powassan virus inside vero cells |  | Hermance et al., 2017 |
| miR-8 | Development, Reproduction | *SWIM* | Bryant et al., 2010; Feng et al., 2018; Lucas et al.,2015 |
| miR-12 | critical for persistence of wolbachia | *DNA replication licensing (MCM6), monocarboxylate transporter (MCT1)* | Osei-Amo et al., 2012 |
| miR-263a | Development |  | Hu et al., 2015 |

miR-8 is upregulated in *E. chaffeensis*-infected tick salivary glands. Its role in innate immunity is unknown so far, but our results have indicated its possible role against Ech-infection, but further work is required for validation. It is a conserved miRNA, and previous studies have shown its role in development and reproduction. Its expression was found up-regulated in *Aedes aegypti* during the pupation stage, with its highest expression levels observed in the mid-pupal period. Previous work by Bryant et al. (2010) revealed the upregulation of miR-8 in the fat body of blood-fed female mosquitoes, suggesting a potential regulatory function in the reproduction of Ae. aegypti. In contrast to *Ae. aegypti*, miR-8 shows abundant expression at different developmental stages of *Anopheles stephensi* (Feng et al., 2018) and is equally expressed in uninfected and infected *Ae. albopictus* saliva following CHIKV infection (Maharaj et al., 2015).

miR-263a was also found upregulated in *E. chaffeensis* infected tick salivary glands indicating its possible role in either innate immunity or transmission, which needs to be revealed by further work. Previous studies have shown its role in development. It was found highly expressed in uninfected and infected *Ae*. *aegypti* saliva (Maharaj et.al. 2015), and is amongst the most highly expressed miRNAs across developmental stages in many mosquito species (Hu et.al.,2015).

miR-12 was found downregulated in *E. chaffeensis* infected tick salivary glands indicating its possible role in activating immune pathways, and its downregulation might have a probable effect on *Ehrlichia* survival, and therefore successful transmission. Further work is required to validate this hypothesis. Although its immunological role is unknown in previous studies. Although previous work has shown its role in affecting *Wolbachia* density in mosquitoes (Osei-Amo et.al. 2012). The preferential expression of miR-12 in *Anopheles gambiae* occurs in the thorax of both males and females, predominantly in midguts and twice as much in their heads (Winter F et.al.,2007), and its targets are DNA replication licensing factor (MCM6) and monocarboxylate transporter (MCT1) genes, as validated in *A.aegypti*, by which it affects *Wolbachia* density in host cells (Osei-Amo et.al. 2012). Our in silico data indicates that miR-279 is upregulated in the salivary glands of ticks infected with Ehrlichia chaffeensis, suggesting it may have a role in the pathogen’s survival or its transmission. These hypotheses, still in the realm of speculation, warrant further study to be substantiated. Additionally, a recent study has proposed that miR-279 might influence the resistance to the insecticide deltamethrin. It does this by regulating the expression of its target gene CYP325BB1, which codes for the enzyme cytochrome P450 325bb1, in the mosquito species *Culex pipiens* pallens (Li et al., 2021). Given that the functions of microRNAs are conserved, this research work might offer substantial insights into the mechanisms driving acaricide resistance, which would be essential in developing new and effective strategies for tick control in the future.

### Validation of in silico differentially expressed microRNAs by qRT-PCR

The expression levels of differentially expressed miRNAs were validated using qRT-PCR assays on *E. chaffeensis*-infected and uninfected tick tissues (Figure 5). The qRT-PCR patterns of the differentially expressed miRNAs were consistent with the next-generation sequencing (NGS) results for the majority of evaluated miRNAs. However, inconsistencies between the NGS and qRT-PCR data patterns were detected. These discrepancies may arise from the different methodologies used to quantify miRNA expression (Saldana et al., 2017).

**Figure 5.**
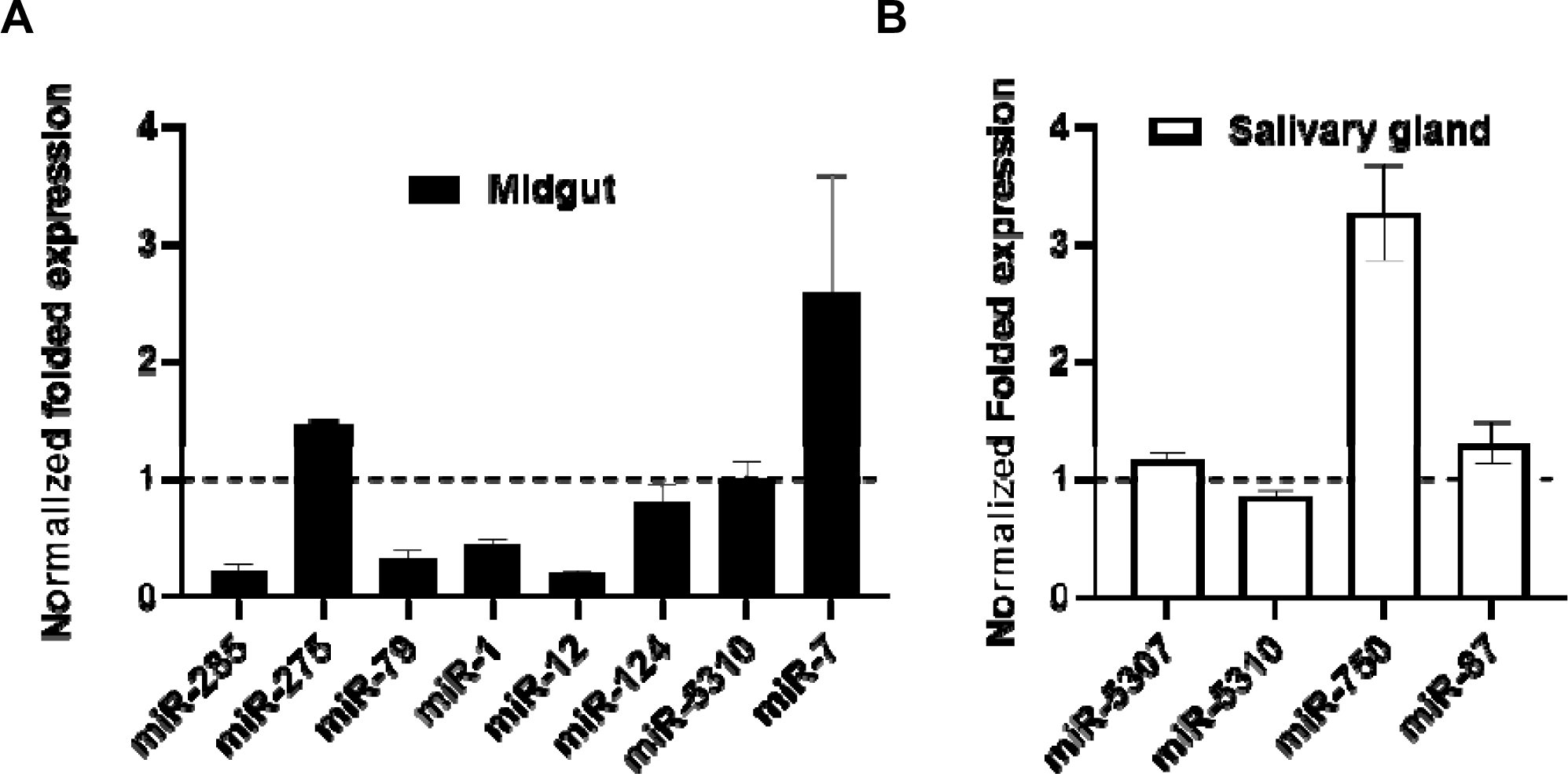
qPCR validation of differentially expressed miRNAs (in silico)in *Ehrlichia chaffeensis* infected and partially fed tick tissues. A) Midgut B) Salivary glands. Expression of miRNAs was normalized with clean and partially fed tick tissues (indicated as 1 on the y-axis). Statistical significance for qRT-PCR-based differential expression was determined by the 2-tailed Student’s t-test where * is *p*<0.05. At least three biological replicates were used in each of the experiments.

### miRNA inhibition in tick tissues reduced *E. chaffeensis* load

Three differentially expressed miRNAs Aam-miR-87, Aam-miR-750, and Aam-miR-275 were selected for the miRNA inhibition assay in tick tissues based on their putative role in tick immune responses (Table 1). As depicted in figures 2 and 3, Aam-miR-87 and Aam-miR-750 were found to be upregulated in the salivary glands of *Ehrlichia chaffeensis*-infected ticks, while Aam-miR-275 was upregulated in the midgut of ticks infected with *Ehrlichia chaffeensis*. The results of our miRNA inhibitory experiments revealed that suppressing these microRNAs individually reduced the *E. chaffeensis* load in tick tissues suggesting a role for miRNAs in tick immunity (Figure 6), although further validation is necessary. Subsequent studies will explore the targets of these microRNAs to gain a deeper understanding of their importance in pathways essential for the survival *of E. chaffeensis*. Additional functional studies will examine how miRNAs and their specific targets impact pathways that affect tick vector competence. Further investigation is also needed to explore additional differentially expressed miRNAs identified in this study, which could potentially offer valuable insights for preventing *E. chaffeensis* infection.

**Figure 6.**
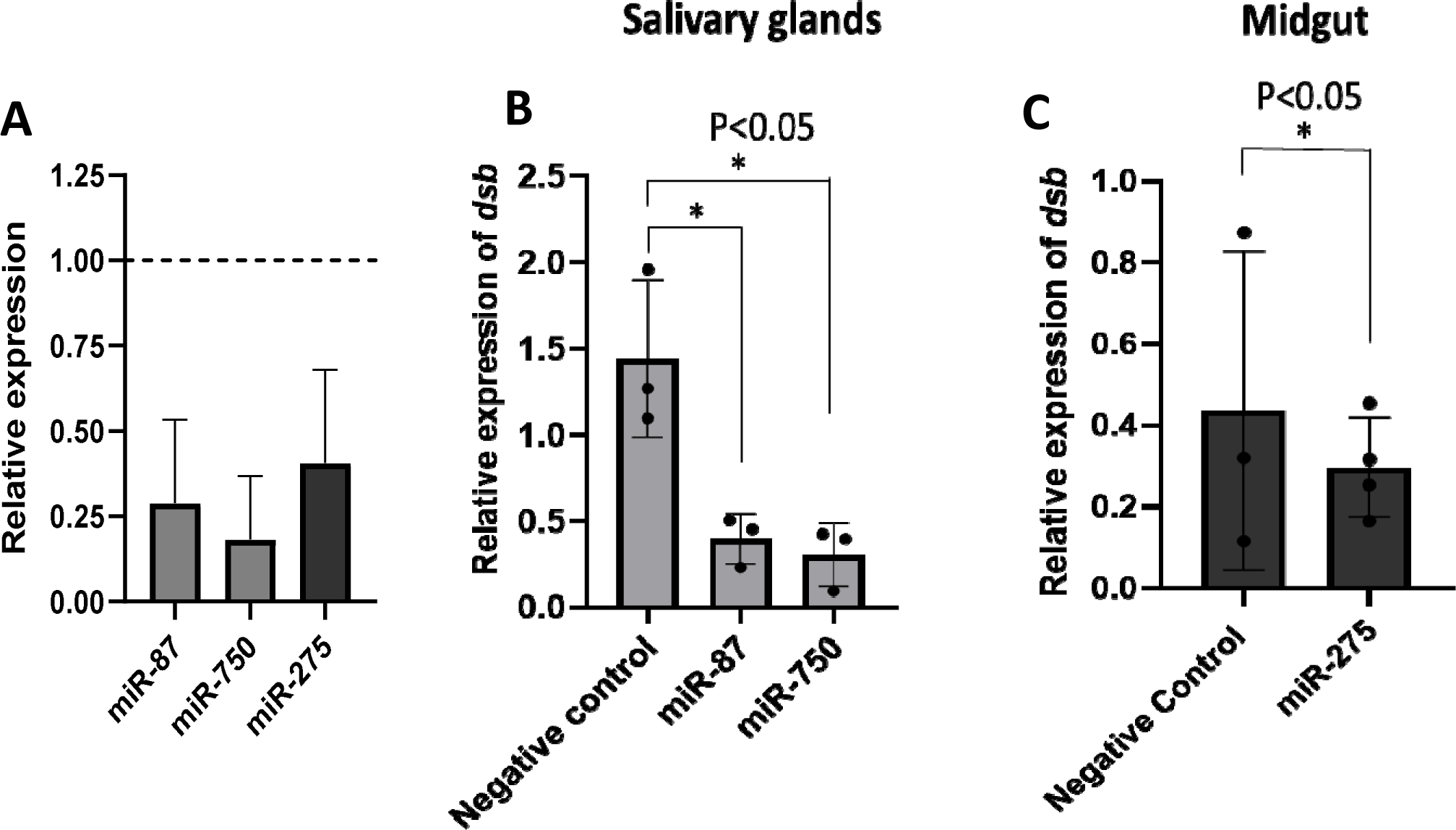
microRNA inhibition reduces *Ehrlichia chaffeensis* load in tick tissues. A) microRNA inhibition in tick tissues (MG and SG). miR-87 and miR-750 were inhibited by ∼75% and ∼80% in tick salivary glands while miR-275 was inhibited by ∼63% in tick midgutExpression of microRNAs in negative control tick tissues were given a normalized fold expression value of 1, as represented by the dashed line. Tick microRNA inhibition resulted in E. chaffeensis load reduction (dsb gene) in B) tick salivary glands and C) midguts. At least three biological replicates were used in each of the experiments. Statistically significant change (P<0.05) is indicated by an asterisk (*).

### Limitations and conclusion

This study highlights the differential expression of miRNAs in *E. chaffeensis-*infected tick tissues, which could significantly influence the survival, colonization, and transmission of *E. chaffeensis*. Additionally, these miRNAs may play a role in the tick’s immune response against the pathogen. Selected microRNAs miR-87, miR-750, and miR-275 have shown promising results against *E. chaffeensis* survival or colonization in ticks. Further investigation of other differentially expressed miRNAs through miRNA inhibitory experiments is needed to explore these aspects. These tick-specific and *E. chaffeensis* specific differentially expressed miRNAs could provide potent avenues for treating or inhibiting *E. chaffeensis*.

Here it is noteworthy to mention that analyzing microRNA sequencing data (from Illumina i.e. short reads) within the context of a long-read sequenced genome may pose challenges in read mapping, alignment, and data integration due to the differences in read lengths and sequencing technologies. Aligning short Illumina reads to a long-read sequenced genome could result in decreased mapping efficiency, especially in regions with structural variations or repetitive elements. This may impact the accuracy of microRNA expression profiling and annotation. While long-read sequences may better capture genomic heterogeneity and structural variations compared to short reads, and provide valuable insights into genome architecture, they may also introduce challenges in accurately identifying and characterizing microRNAs, particularly in regions of high complexity or variation. Long-read sequencing technologies may have significantly higher error rates, which can pose challenges in accurately reconstructing the genome, mapping microRNA sequences, and distinguishing true variations from sequencing errors.

## Data availability statement

The raw small RNA sequences were deposited into the NCBI Sequence Read Archive (SRA) repository under the BioProject ID PRJNA992656.

## Ethics statement

The animal study was reviewed and approved by the University of Southern Mississippi’s Institutional Animal Care and Use Committee (IACUC protocols # 15101501.3 and 17101206.2). The study was conducted in accordance with the local legislation and institutional requirements.

## Author contributions

DK: Conceptualization, Formal analysis, Investigation, Methodology, Writing-original draft, Writing-review & editing. KRB: Investigation, Methodology, Resources, Writing-review & editing. YR: Resources, review & editing. SK: Conceptualization, Funding acquisition, Investigation, Project administration, Resources, Supervision, Writing-Original draft, Writing-review & editing.

## Acknowledgment

This work utilized the Magnolia High-Performance Computing at USM (http://magnolia.usm.edu).

